# *Aspergillus*-mediated allergic airway inflammation is triggered by dendritic cell recognition of a defined spore morphotype, a process that can be targeted via antifungal therapeutics

**DOI:** 10.1101/2024.01.11.575032

**Authors:** EL Houlder, S Gago, G Vere, D Conn, S Khan, D Thomson, MW Shepherd, R Lebedinec, GD Brown, M Bromley, AS MacDonald, PC Cook

## Abstract

**Background:** Exposure to fungi, especially *Aspergillus fumigatus (A.f.)*, can elicit potent allergic inflammation that triggers and worsens asthmatic disease. Dendritic cells (DCs), initiate allergic inflammatory responses to allergic stimuli. However, it is unclear if *A.f.* spores during isotropic growth (early spore swelling) can activate DCs to initiate allergic responses or if germination is required. This lack of basic understanding of how *A.f.* causes disease is a barrier to the development of new treatments.

**Objective:** To show that a precise *A*.*f*. morphotype stage during spore swelling can trigger DCs to mediate allergic inflammatory responses and ascertain if antifungal therapeutics can be effective at suppressing this process.

**Methods:** We employed an *A.f.* strain deficient in pyrimidine biosynthesis (ΔpyrG) to generate populations of *A.f.* spores arrested at different stages of isotropic growth (swelling) via temporal removal of uracil and uridine from growth media. These arrested spore stages were cultured with bone marrow derived DCs (BMDCs), and their activation measured via flow cytometry and ELISA to interrogate which growth stage was able to activate BMDCs. These BMDCs were then adoptively transferred into the airways, to assess if they were able to mediate allergic inflammation in naive recipient mice. Allergic airway inflammation *in vivo* was determined via flow cytometry, ELISA and qPCR. This system was also used to determine if antifungal drug (itraconazole) treatment could alter early stages of spore swelling and therefore BMDC activation and *in vivo* allergic inflammation upon adoptive transfer.

**Results:** We found that *A*.*f*. isotropic growth is essential to trigger BMDC activation and mediate allergic airway inflammation. Furthermore, using time arrested *A.f.* stages, we found that least 3h in growth media enabled spores to swell sufficiently to activate BMDCs to elicit allergic airway inflammation *in vivo*. Incubation of germinating *A.f.* with itraconazole reduced spore swelling and partially reduced their ability to activate BMDCs to elicit *in vivo* allergic airway inflammation.

**Conclusion:** In summary, our results have pinpointed the precise stage of *A*.*f*. development when germinating spores are able to activate DCs to mediate downstream allergic airway inflammation. Furthermore, we have identified that antifungal therapeutics can be effective in reducing the potential of *A.f.* spores to stimulate allergic responses, highlighting a potential mechanism by which antifungal treatment might help to prevent the development of fungal allergy.

## Introduction

Fungi are abundant in the environment and are a major driver of asthma (1,2). At least 10 million people worldwide display significant sensitivity to the mould *Aspergillus fumigatus* (*A.f.*), which increases asthma severity (3) and can lead to the development of debilitating chronic fungal infections (4,5) such as allergic bronchopulmonary aspergillosis (ABPA) and severe asthma with fungal sensitisation (SAFS) (6). Despite the significant clinical burden of these diseases, treatment options are limited. Moreover, there is much debate on whether it is preferable to administer therapeutics, such as corticosteroids, that dampen the underlying allergic inflammation but may interfere with antifungal immunity (7) or utilise antifungal therapies, such as the drug itraconazole, which have been shown to improve health for individuals with chronic allergic fungal conditions (8,9). Although the precise mechanism(s) that might explain how such treatments are effective are unknown, these observations suggest that *A.f.* plays a critical, yet underappreciated, role in mediating responses that underpin allergic airway inflammation.

The immune events that cause individuals to become sensitised to *A.f.* spores and develop allergic airway inflammatory disease is unclear. In healthy settings, the majority of spores are removed in the upper airways, with their small size (2 – 3 μM) allowing around 20% of inhaled *A.f.* spores to reach the alveoli where they are thought to be cleared by cells of the epithelial barrier, along with innate immune cells such as macrophages and dendritic cells (DCs) (10–13). However, relatively few studies have focused on whether *A.f.* spores activate these cells to co-ordinate allergic inflammatory responses. In the lung, DCs sample the airway and migrate to the draining lymph nodes (dLNs) to mediate antigen specific T cell responses against a range of stimuli, including fungi (14–16). We have recently identified that DCs orchestrate CD4^+^ T cell responses to repeated exposures of *A.f.* spores to generate a mixed type 2 (eosinophilia with IL-4, IL-5 and IL-13) and type 17 (neutrophilia with IL-17) inflammatory cytokine environment (17). These immune changes mediate the hallmark symptoms of asthma, including airway hypersensitivity, luminal narrowing, smooth muscle hyperplasia and mucus over production (18). In contrast to other allergens, such as house dust mite (HDM), little is known about whether *A.f.* spores actively reveal factors that trigger DC mediated sensitisation and their promotion of fungal allergic airway inflammatory disease.

If not rapidly cleared from the airways, within hours *A.f.* spores can undergo morphogenetic changes as they swell and then germinate to form hyphae (19,20). Resting *A.f.* spores are coated in an immunologically inert dense α-1,3-glucan, melanin and rodlet hydrophobin layer (21–26). This layer is shed as spores break dormancy and undergo isotropic growth (swelling) (21) exposing pathogen associated molecular patterns (PAMPs), such as carbohydrate motifs including β-1,3-glucan, that can be recognised by innate immune cells and trigger potent inflammatory responses (26–29). Furthermore, numerous *A.f.* allergens have been identified, with hyphal proteases (Asp f 5, Alp1 and Asp f 13) particularly implicated in mediating allergic inflammatory responses (30–33). However, whether these allergens are expressed in early germinating spores and mediate initial sensitisation via DCs is unknown. Deeper understanding of the potential immunogenicity of molecules exposed on the swollen spores (34) is vital to precisely map when *A.f.* spores develop allergenic properties, and to identify how this process may be therapeutically manipulated (Ortiz et al., 2019).

In this study, we have demonstrated an absolute requirement for *A.f.* spore swelling to induce immune activation, both *in vitro* and *in vivo*, with a particular focus on DCs due to their crucial role in directing allergic airway inflammation. Further, we have identified the stage in *A.f* isotropic growth at which spores are able to activate BMDCs to initiate allergic airway inflammation. Finally, we have shown that treatment with the antifungal itraconazole can reduce the allergenicity of *A.f.* spores, and therefore their capacity to activate BMDCs to initiate allergic airway inflammation. These results not only precisely define the development of allergenicity during *A.f.* spore swelling but also provide mechanistic evidence to explain the therapeutic benefit of antifungal treatment in allergic airway inflammation.

## Methods

### *Aspergillus* strains and culture

*A.f* A1160 *ΔpyrG (pyrG*^−^) was used to generate stage arrested *A.f.* spores, while A1160 *pyrG* (referred to as A1160 WT) strain was used as control for comparisons for *in vitro* and i*n vivo A.f.* challenge (35–37). The A1160 strain was derived from CEA10, an invasive aspergillosis clinical isolate (38). In experiments involving DC transfer to recipient mice, mice were challenged with spores of the parental CEA10 strain. *A.f.* was grown on Sabouraud Dextrose Agar (SAB, Oxoid), at 37⍰°C supplemented with 5 mM uridine and 5 mM uracil (u/u, both Sigma) for the A1160 *ΔpyrG* strain. Spores were harvested resuspended in RPMI (Sigma) at 1.6 ×10 spores/ml in the presence or absence of 5 mM u/u. In some experiments these were further supplemented with 2 μg/ml itraconazole (Thermo Scientific). Stage arrested spores were generated by incubation at 37⍰°C (with shaking) for defined periods and washed in non-supplemented RPMI (Gibco) to remove excess u/u. To aid counting, spores were sonicated for 3 sec at 20% amplitude to disperse clumps. Swollen spores were stored in non-supplemented RPMI 4⍰°C for up to 24h prior to addition to bone marrow dendritic cell (BMDC) culture or assessed directly by flow cytometry.

### BMDC culture

BMDCs were generated with granulocyte–macrophage colony stimulating factor (GM-CSF) as previously described (39). In brief, 2 × 10 bone marrow cells were seeded in complete medium (RPMI-1640 (Sigma) containing 20⍰ng/ml GM-CSF (Peprotech), 10% foetal calf serum (Sigma), 2⍰mM L-glutamine (Gibco), 50⍰U/ml penicillin and 50⍰μg/ml streptomycin (Life Technologies). Cells were cultured at 37⍰°C in a humidified atmosphere of 5% CO_2_. On day 3, 10⍰ml of complete medium was added and on days 6 and 8, 9⍰ml of media was gently aspirated and replaced with 10⍰ml of fresh complete medium. Following 10 days of culture, BMDCs were harvested and replated at 2 × 10 cells/ml for further assays. *A.f.* spores were added to BMDCs at multiplicity of infection (MOI) 5:1 and incubated for 5h at 37⍰°C in a humidified atmosphere of 5% CO_2_, in complete medium with 5 ng/ml GM-CSF. Media or 250 ng/ml LPS (Sigma)were added as negative and positive controls respectively.

### Murine models

C57BL/6 (Envigo) mice were maintained under specific pathogen free conditions at the University of Manchester (licence number P44492AC9) or University of Exeter (license number P6A6F95B5). All animal studies were ethically reviewed and carried out in accordance with Animals (Scientific Procedures) Act 1986 and the GSK Policy on the Care, Welfare and Treatment of Animals. Female mice aged 6-17 weeks were used for analysis. No *a priori* sample size calculations were performed to determine number of mice used per group. All mice were included in the analysis, with no exclusions. For statistical calculations individual mice were regarded as the experimental unit. Blinding and randomisation of treatment groups was not performed. To account for confounding “cage-effects” mice from different treatment groups were co-housed.

In repeat *A.f.* exposure models, experimental mice were exposed intranasally (*i.n.*) to 4 × 10^5^ *A.f.* spores (A1160) in 30 μl PBS 0.05% Tween or vehicle control on days 0, 2, 4 7, 9 and 11. Mice were euthanised at day 12 post first *A.f.* dose. In DC transfer models, mice were sensitised via *i.n.* transfer of 5 × 10^4^ BMDCs (day 0), challenged with *A.f.* (CEA10) on days 12 and 13 post transfer (4 × 10^5^ each day) and euthanised on day 14.

### Cell isolation

For analysis of cellular changes *in vivo,* cells were isolated from the bronchiolar lavage (BAL), lung tissue, and mediastinal lymph nodes. BAL was collected by washing the lungs with PBS containing 2% FBS and 2 mM EDTA (Sigma). Lungs were processed via incubation at 37 °C with 0.8 U/ml Liberase TL (Sigma) and 80 U/ml DNase I type IV (Sigma) in HBSS (Gibco), as previously described (39). After 40 min the digestion was halted with PBS containing 2% FBS and 2 mM EDTA, and the suspension passed through a 70 μm cell strainer. Red blood cells (RBCs) were lysed using RBC lysis buffer (Sigma). To assess cytokine secretion potential, lung cells were stimulated *ex vivo* for 3h at 37°C with 20 ng/ml PMA (Sigma) and 1 µl/ml GolgiStop (BD) in X-vivo-15 (Lonza), supplemented with 1% L-glutamine (Gibco) and 0.1% β-mercaptoethanol (Sigma). Cultures were carried out in 96 well round-bottom plates, 4 × 10^5^ cells per well in 200 µl final volume.

### Flow cytometry

Isolated cell suspensions were washed with PBS and stained for viability with ZombieUV (1:2000; Biolegend). Samples were then blocked with 5 μg/ml αCD16/CD32 (2.4G2; BioLegend) in FACS buffer (PBS containing 2% FBS and 2 mM EDTA) before staining for surface markers at 4°C for 30 min. Post staining, cells were washed twice in FACS buffer and then fixed in 1% paraformaldehyde in PBS for 10 min at room temperature. For detection of intracellular antigens, cells were fixed with BD cytofix/cytoperm (BD) for 1h. Cells were then washed three times with eBioscience permeabilization buffer (Thermofisher) before overnight staining with intracellular cytokine antibodies in eBioscience permeabilization buffer (ThermoFisher). Samples were acquired on a BD Fortessa with FacsDiva (BD) software and analysed with Flowjo v10 (Tree Star). Gating was informed by the use of fluorescence minus one controls. Flow cytometry gating schemes are shown in Supplementary Figures 1-3.

### ELISA

ELISAs were performed on BAL and BMDC culture supernatants using paired mAbs and recombinant cytokine standards, or DuoSets following manufacturer’s instructions (BioLegend, eBio-science, BD, R&D Systems, and PeproTech).

### RNA extraction and qPCR gene expression

Lung tissue was collected in RNAlater (ThermoFisher) for storage at −80°C prior to subsequent processing. Once thawed, tissues were homogenised in RLT lysis buffer in a Geno/Grinder (SPEX) and RNA isolated using RNeasy columns (Qiagen) following manufacture instructions. Complementary DNA was generated from extracted RNA using SuperScript-III and Oligo-dT (ThermoFisher). Relative quantification of genes of interest was performed by qPCR analysis using QuantStudio Pro 7 system (ThermoFisher) and SYBR Green master mix (ThermoFisher), compared with a serially diluted standard of pooled cDNA. Expression was normalised to GAPDH (17).

### Statistical analysis

For all figures, to compare between groups, mixed linear models were fitted utilising Rstudio (R Core Team) and the lme4 package (40). To account for random variation, experimental day was designated a random effect variable, with timepoint or genotype as fixed effect variables. To compare multiple groups a post-hoc Tukeys HSD test was used using the multcomp package (41). For Supplementary Figure 5 GraphPad Prism 8.0 was used, with multiple comparisons testing applied after fitting a mixed model using a compound symmetry covariance matrix and Restricted Maximum Likelihood (REML). Bar charts show mean and standard error of the mean (SEM).

## Results

### *A.f.* spore viability is required for induction of fungal allergic airway inflammation

To determine the requirement for *A.f.* viability in mediating early immune sensitisation events, we utilised a murine model of fungal allergic airway inflammation (17,42) which involved administering repeated doses (4 × 10) of live or PFA-killed spores via *i.n.* transfer (Fig. 1A). As expected, 24h after 6x *A.f.* doses (d12), we found that mice which received live spores had a significant influx of granulocytes and lymphoid cells in the lung airway (via analysis of cells isolated from the BAL fluid) (Fig. 1B). This was also accompanied by markedly increased type 2 (expressing IL-4, IL-5 and IL-13) and type 17 (expressing IL-17) CD4^+^ T cells in the lung (Fig. 1C), confirming previous reports that repeat *i.n.* dosing with live *A.f.* spores induces hallmarks of allergic airway inflammation (42). Strikingly, we found that the response in mice that received PFA-killed spores was significantly reduced, and similar to PBS controls (Fig. 1B&C), confirming and extending previous observations (42), that spore viability is crucial to induce allergic airway inflammation.

**Figure 1.**
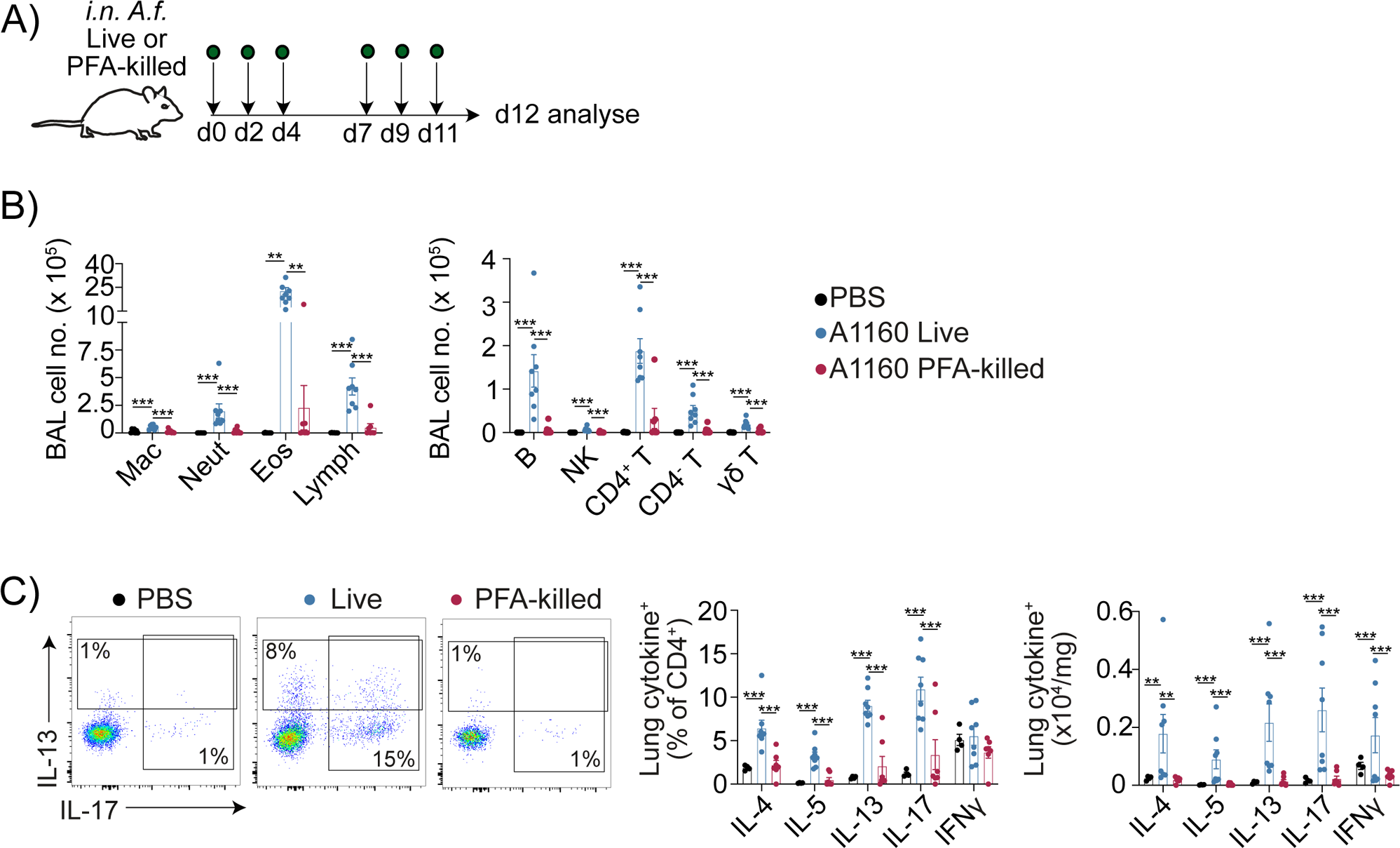
PFA-killed spores are unable to induce fungal allergic airway inflammation. **A**) C57BL/6 mice were exposed to six doses of live spores (A1160), PFA killed spores (A1160) (0.4×10^6^ per dose) or PBS *i.n.* on the indicated days and tissues harvested 24h after the sixth dose (d12). **B**) The numbers of different cell populations isolated from BAL fluid of *A.f.* or PBS exposed mice were assessed by flow cytometry. **C**) Flow cytometry plots show CD4^+^ T cell intracellular cytokine production following *ex vivo* stimulation with PMA/ionomycin of lung cells from *A.f.* or PBS exposed mice. Data combined from two independent experiments (n = 7-8 per group). Each data point is an individual mouse. Linear mixed effect modelling applied, with experimental repeat as a random effect variable, to compare multiple groups a post-hoc Tukeys HSD test was used. *=P <0.05,**=P <0.01, ***=P <0.001, ****=P <0.0001.

A major drawback when using PFA to kill spores is that it can dramatically alter cell wall surface epitopes (43), as well as halt spore metabolism, both of which could influence the ability of spores to interact with and activate DCs during allergic airway inflammation. Therefore, we utilised the Δ*pyrG A.f.* strain (lacking orotidine-51-monophosphate decarboxylase, an essential enzyme in the pyrimidine biosynthetic pathway), which is viable but unable to develop due to disrupted uridine synthesis to investigate whether spore swelling is crucial for *A.f.* induction of allergic airway inflammation *in vivo* (35–37). Mice that were repeatedly exposed to non-swollen *ΔpyrG* spores showed significantly reduced myeloid, granulocyte and lymphoid cell populations in the BAL fluid, compared to mice exposed to WT (*pyrG^+^*) spores (Fig. 2A & B). Furthermore, non-swollen Δ*pyrG* spores triggered reduced secretion of mediators associated with allergic inflammation in the airway (measuring RELMα and chitinase like protein (Chi3l3) in the BAL fluid) compared to WT spores, which can germinate in the host (Fig. 2C). Surprisingly, despite these differences in inflammatory cell types, non-swelling Δ*pyrG* spores induced a similar proportion of CD4^+^ T cells to produce type 2 and type 17 cytokines (Fig. 2D), although there was a trend for reduced numbers of these cells. However, we found expression of genes associated with type 2 inflammation (*Retnla*, *Chi3l3*, *Il5*) were significantly reduced in the lung tissue of mice that received non-swelling Δ*pyrG* versus WT spores (Fig. 2E). In summary, these results show that live spores that are incapable of isotropic growth have a much-reduced capacity to trigger fungal allergic airway inflammation.

**Figure 2:**
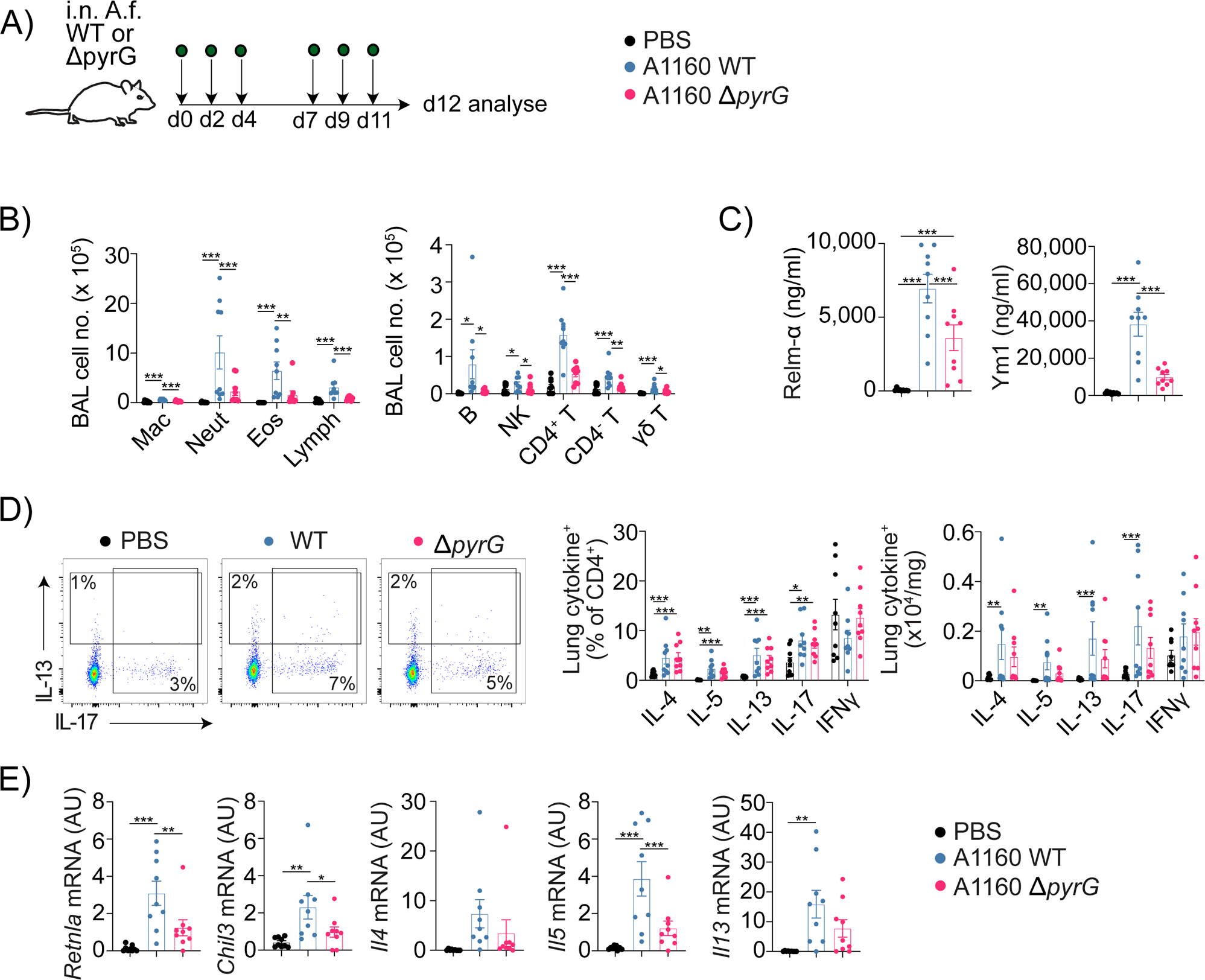
Fungal allergic airway inflammation in response to *A.f.* is dramatically reduced when spore swelling is arrested. **A**) C57BL/6 mice were exposed to six doses of *A.f.* spores (A1160), non-germinating Δ*pyrG A.f.* spores (0.4×10^6^ per dose) or PBS *i.n.* on the indicated days and tissues were harvested 24h after the sixth dose (d12). **B**) Immune cell populations in the BAL fluid were assessed by flow cytometry. **C**) Secretory factors in the BAL fluid were quantified by ELISA. **D**) Representative flow cytometry plots and graphs show lung CD4^+^ T cell intracellular cytokine staining after *ex vivo* stimulation with PMA/ionomycin. **E**) Lung tissue mRNA expression was assessed by qPCR (normalised against *Gapdh*, a.u.). Data combined from two independent experiments (n = 9 per group). Each data point is an individual mouse. Linear mixed effect modelling applied, with experimental repeat as a random effect variable, to compare multiple groups a post-hoc Tukeys HSD test was used. *=P <0.05,**=P <0.01, ***=P <0.001, ****=P<0.0001.

### At least 3h of isotropic growth is essential for spores to elicit BMDCs to mediate fungal allergic airway inflammation

After identifying that spore swelling was essential for *in vivo A.f.* induction of allergic airway inflammation (Fig. 2), we wanted to define the precise timepoint at which spores were able to activate BMDCs to mediate allergic inflammation. As *A.f.* spores are rapidly cleared after inhalation, we suggest that immune responses to early stages of spore swelling may be critical. To identify at which point after breaking dormancy A.f spores were able to elicit an immune response, we further utilised *ΔpyrG* spores with timed addition and then removal of uridine and uracil (u/u) supplemented media, to arrest isotropic growth at defined timepoints (Fig. 3A). We confirmed this was achieved by measuring spore morphogenesis via flow cytometry and microscopy, observing that spore size began to increase at 3h of culture, with a significant increase in the proportion of swollen spores at 4h (Fig. 3B and Supplementary Fig. 4A). Arrested spores remained viable after the removal of u/u, and they were able to germinate when u/u was re-introduced 24h later (Supplementary Fig. 4B). These stages of growth-arrested *ΔpyrG* spores were then each separately cultured with BMDCs to determine their ability to trigger DC activation (Fig. 3C). We consistently found that 3h of swelling was the earliest isotropic growth stage that triggered significant BMDC activation, indicated by increased surface expression of costimulatory molecules (CD86, CD80 and MHCII) alongside secretion of pro-inflammatory cytokines (IL-6, IL-12p40 and TNFα) (Fig. 3D & E). As expected, later growth stages (4h & 6h) induced the most marked BMDC activation (Fig. 3D & E). In comparison all spores grown in the absence of u/u, 0h, 1h or 2h spore stages did not trigger BMDC activation (Fig. 3D & 3E). Our findings using the arrested *ΔpyrG* system were later confirmed using PFA-killed spores, pre-swollen for 4h, which were able to induce immune activation in BMDCs (Supplementary Fig. 5). Importantly, incubation of BMDCs with live spores for 5h only induced minor BMDC immune activation likely due to the fact that more immunogenic stages (3h+) had reduced time to interact with the BMDCs (Supplementary Fig. 5). This highlights the importance of using a growth arrested *A*.*f*. to allow sufficient time for immune activation by each spore morphotype. Taken together, utilising the A1160 *ΔpyrG* strain and a novel approach of timed addition and removal of u/u, we have pinpointed the earliest morphotype spore stage that is able to trigger BMDC activation.

**Figure 3:**
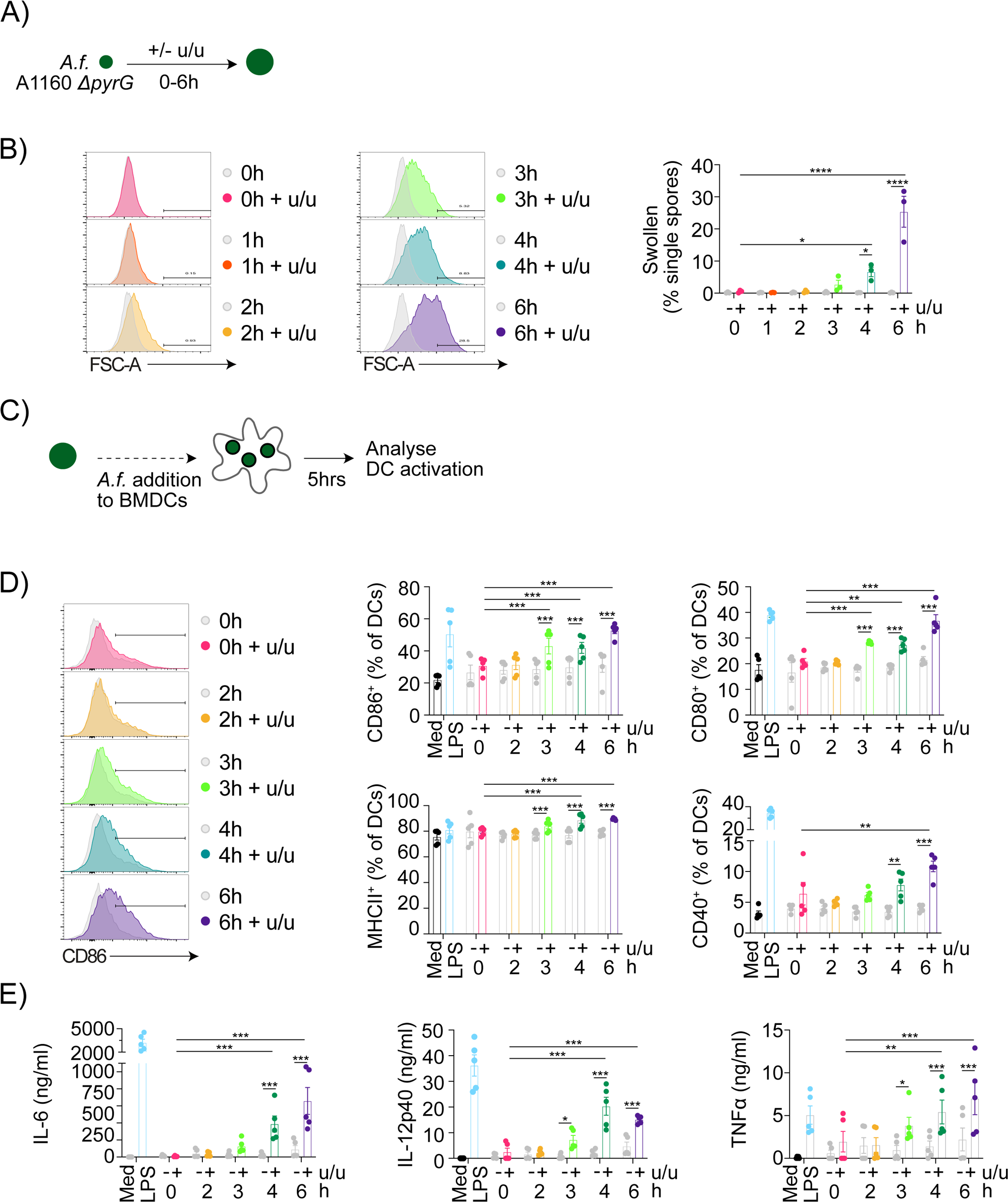
*A.f.* activation of DCs requires spores to undergo at least three hours of isotropic growth. **A**) *ΔpyrG* spores were incubated in the presence or absence of uridine and uracil (u/u) with growth arrested by the removal of u/u media at the indicated timings (between 0 - 6h). **B**) Spore size was determined by flow cytometry. **C**) Growth arrested *ΔpyrG* spores were incubated with DCs for 5h (MOI 5:1) and DC activation analysed. **D)** DC expression of co-stimulatory molecules and MHCII was analysed via flow cytometry. **E**) DC cytokine secretion was measured via ELISA. Data combined from two independent experiments (n=5 per group). Each data point is an individual DC culture. Linear mixed effect modelling applied, with experimental repeat as a random effect variable, to compare multiple groups a post-hoc Tukeys HSD test was used. Statistical comparisons to LPS positive control are not shown. *=P <0.05,**=P <0.01, ***=P <0.001, ****=P<0.0001.

Having shown that 3h of *A*.*f*. isotropic growth was sufficient to induce BMDC activation *in vitro*, we wanted to ascertain if these were able to mediate allergic airway inflammation *in vivo*. To test this, we adapted a BMDC adoptive transfer model that we have used previously (39), whereby BMDCs previously exposed to 0h, 2h or 3h arrested *ΔpyrG* spores were administered *i.n.* to naïve recipient mice, followed by two further *i.n.* challenge doses of live *A.f.* spores (days 11 and 12 after BMDC sensitisation) (Fig. 4A). Only mice that were sensitised to BMDCs pulsed with 3h spores displayed increased cell recruitment to the airways, with significantly increased granulocytes (neutrophils and eosinophils) and lymphocyte (NK cells, CD4^+^ & CD8^+^ T cells and γδ T cells) populations in the BAL fluid, when compared to mice sensitised with BMDCs that had been cultured in media alone (Fig. 4B). In addition, there was also a significant increase in the number of activated CD4^+^ T cells (CD44^+^) Fig. 4C), as well as secretory factors associated with allergic inflammation (RELMa and Ym1) in the BAL fluid (Fig. 4D). To confirm these features were reflective of differing capabilities in mediating fungal allergic airway inflammation, we found that adoptively transferred BMDCs that had been cultured with 3h spores were able to induce significant expansion of type 2 cytokine-expressing CD4^+^ T cells (Fig. 4E). In contrast, mice that received BMDCs that had been exposed to 0h or 2h spores showed no evidence of fungal allergic inflammation (Fig. 4). In summary, these data show that BMDC induction of fungal allergic airway inflammation against *A.f.* is dependent on their recognition of a defined spore morphotype that requires at least 3h of growth to become immunogenic.

**Figure 4:**
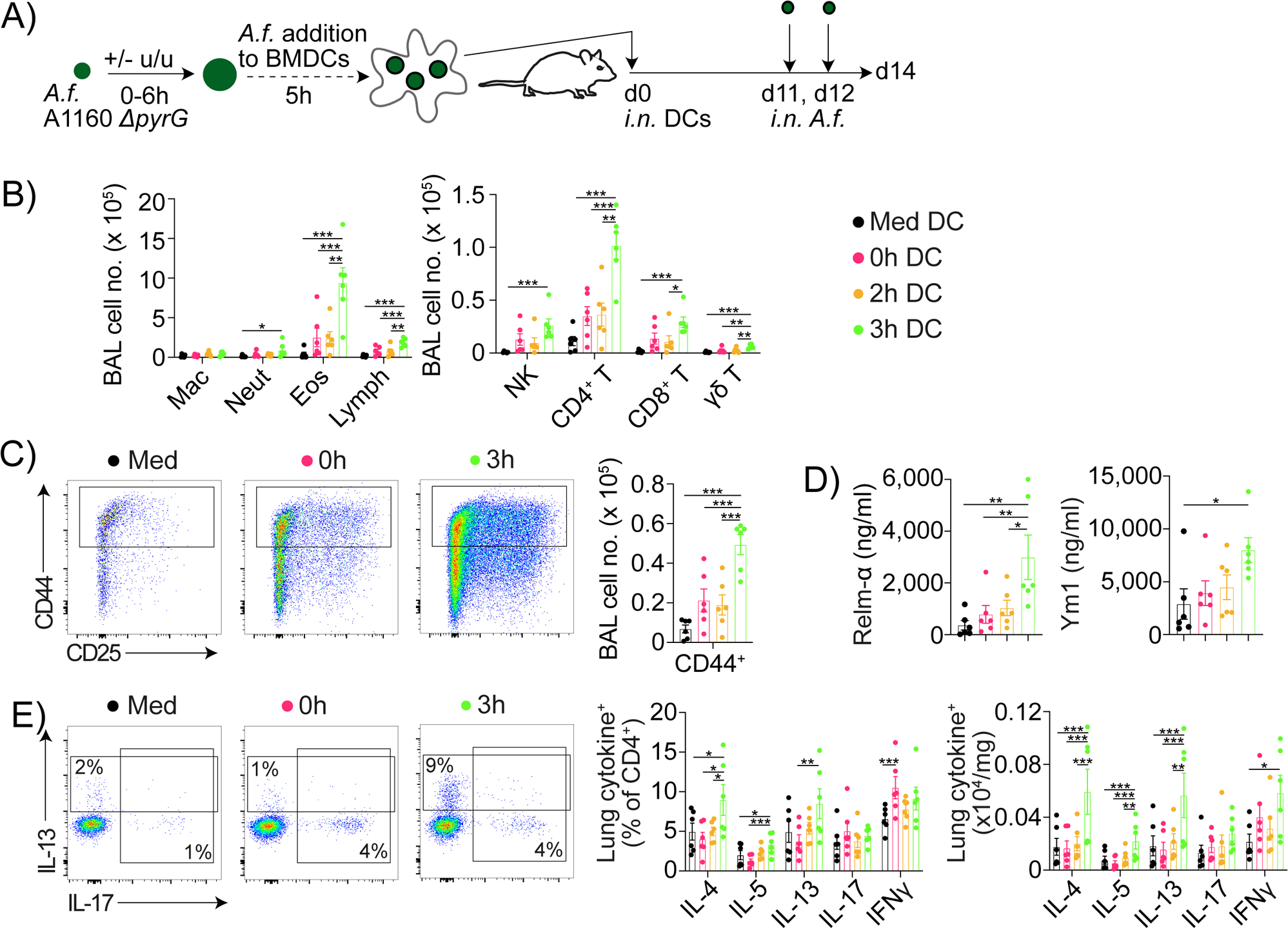
Three hours of isotropic growth is essential for spores to elicit DC ability to mediate fungal allergic airway inflammation. **A**) Arrested *ΔpyrG* spore stages were generated by incubating in the presence of uridine and uracil (u/u), then removed and washed after 0, 2 or 3h. These were then cultured with BMDCs for 5h, prior to *i.n.* transfer of 50,000 DCs into C57BL/6 mice. Mice were challenged *i.n.* at d11 and 12 with 0.4 × 10 live CEA10 *A.f.* spores. **B**) Immune cell populations in the BAL fluid were assessed by flow cytometry. The number of different cell populations isolated from BAL fluid of DC sensitised mice were assessed by flow cytometry. **C**) Representative flow cytometry plots of CD44^+^CD4^+^ T cells in BAL fluid. **D**) Secretory factors in BAL fluid were quantified by ELISA. **E**) Representative flow cytometry plots and graphs show lung CD4 T cell intracellular cytokine staining after *ex vivo* stimulation with PMA/ionomycin. Data combined from two independent experiments (n=6 per group). Each data point is an individual mouse. Linear mixed effect modelling applied, with experimental repeat as a random effect variable, to compare multiple groups a post-hoc Tukeys HSD test was used. *=P <0.05,**=P <0.01, ***=P <0.001, ****=P<0.0001.

### Treatment of *A.f*. with antifungal agents reduces spore capacity to activate BMDC ability to induce allergic airway inflammation

Although it has been reported that antifungal therapies (including azole drugs such as itraconazole) can be a successful treatment strategy for patients with severe fungal asthma (45–47), the mechanisms that underpin this are poorly understood. Therefore, we reasoned that one partial explanation may be that antifungal treatment could interfere with early spore swelling and so reduce their capability to activate BMDCs to induce allergic airway inflammation. To our knowledge, few studies have assessed how antifungal drugs impact early stages of *A.f.* spore swelling. We found that incubating *ΔpyrG A.f.* in the presence of itraconazole (and u/u) for 4h significantly reduced spore swelling (Fig. 5A & B). To test whether this impacted *A.f.* immunogenicity spores were grown in the presence or absence of itraconazole, washed extensively to remove itraconazole, and then cultured with BMDC (Fig. 5A-C). Itraconazole treated 4h spores showed significantly impaired ability to activate BMDCs compared to 4h spores that had not been treated with itraconazole, in terms of induction of surface expression of co-stimulatory molecules (CD40, CD80, CD86 and MHCII) and production of pro-inflammatory cytokines (IL-6, IL-12p40 and TNFα) (Fig. 5D & E). However, while itraconazole treated 4h arrested spores were less proficient at activating BMDCs than untreated 4h spores, the response detected was still significantly above the activation state of BMDCs cultured with 0h spores. Together, these data indicate that *A.f.* treatment with azole-based drugs can reduce, but not eliminate, the ability of early swollen spore stages to activate BMDCs *in vitro*.

**Figure 5:**
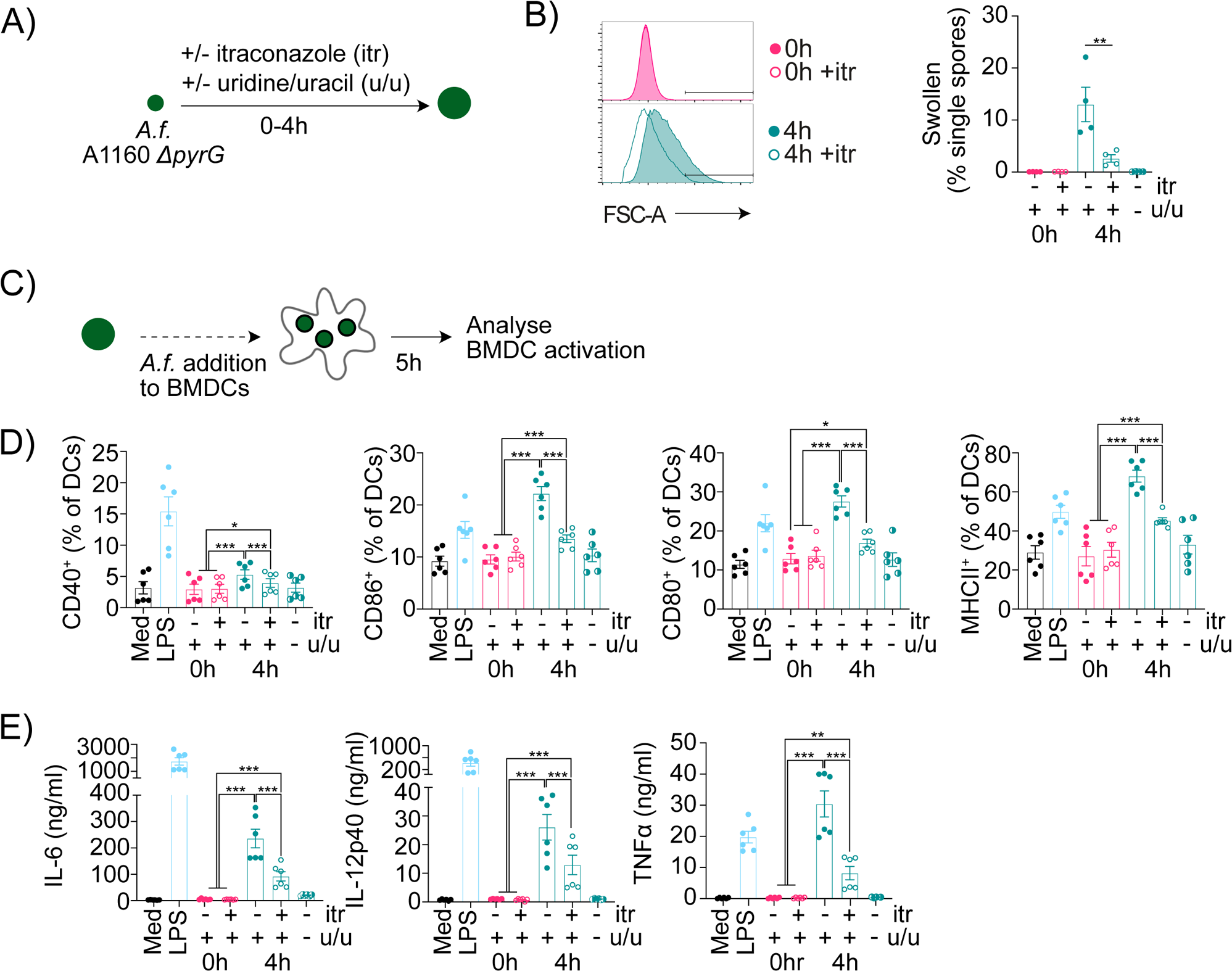
Itraconazole reduces swelling of spores and ability to activate DCs. **A**) *ΔpyrG* spores were incubated with uridine and uracil (u/u) containing media in the presence or absence of itraconazole (itr) then growth was arrested by the removal of u/u media at 0 or 4h. (**B**) Spore size was determined by flow cytometry. **C**) Growth arrested and itraconazole treated *ΔpyrG* spores were incubated with DCs for 5h (MOI 5:1) and DC activation analysed. **D)** DC expression of co-stimulatory molecules and MHCII was analysed via flow cytometry. **E**) DC cytokine secretion was measured via ELISA. Data combined from two independent experiments (n=6 per group). Each data point is an individual DC culture. Linear mixed effect modelling applied, with experimental repeat as a random effect variable, to compare multiple groups a post-hoc Tukeys HSD test was used. Statistical comparisons to LPS positive control are not shown. *=P <0.05,**=P <0.01, ***=P <0.001, ****=P<0.0001.

To test if itraconazole treatment also impacted the ability of early swollen *A.f.* spores to influence BMDC induction of fungal allergic airway inflammation, we transferred BMDCs that had been cultured with 0h or 4h arrested spores germinated in the presence or absence of itraconazole into the airways of naïve recipient mice (Fig. 6A). BMDCs that had been activated with either 4h spores or itraconazole treated 4h spores promoted similar levels of cellular inflammation in the airways of recipient mice (Fig. 6B), with a trend that BMDCs activated with itraconazole treated spores induced less marked eosinophilia. Notably, BMDCs that had been cultured with itraconazole treated spores did display significantly reduced ability to activate CD4^+^ T cells *in vivo*, particularly in terms of number of activated CD44^+^CD4^+^ T cells recruited to the airways (Fig. 6B & C). Analysis of mediators in the BAL fluid revealed that itraconazole treatment of *A.f.* spores did not significantly influence the ability of BMDCs to induce *in vivo* secretion of factors associated with allergic airway inflammation (Fig. 6D). Taken together, these data show that itraconazole treatment of *A.f.* spores disrupts their morphotype transition, while making them less effective at activating BMDCs to induce allergic airway inflammation.

**Figure 6:**
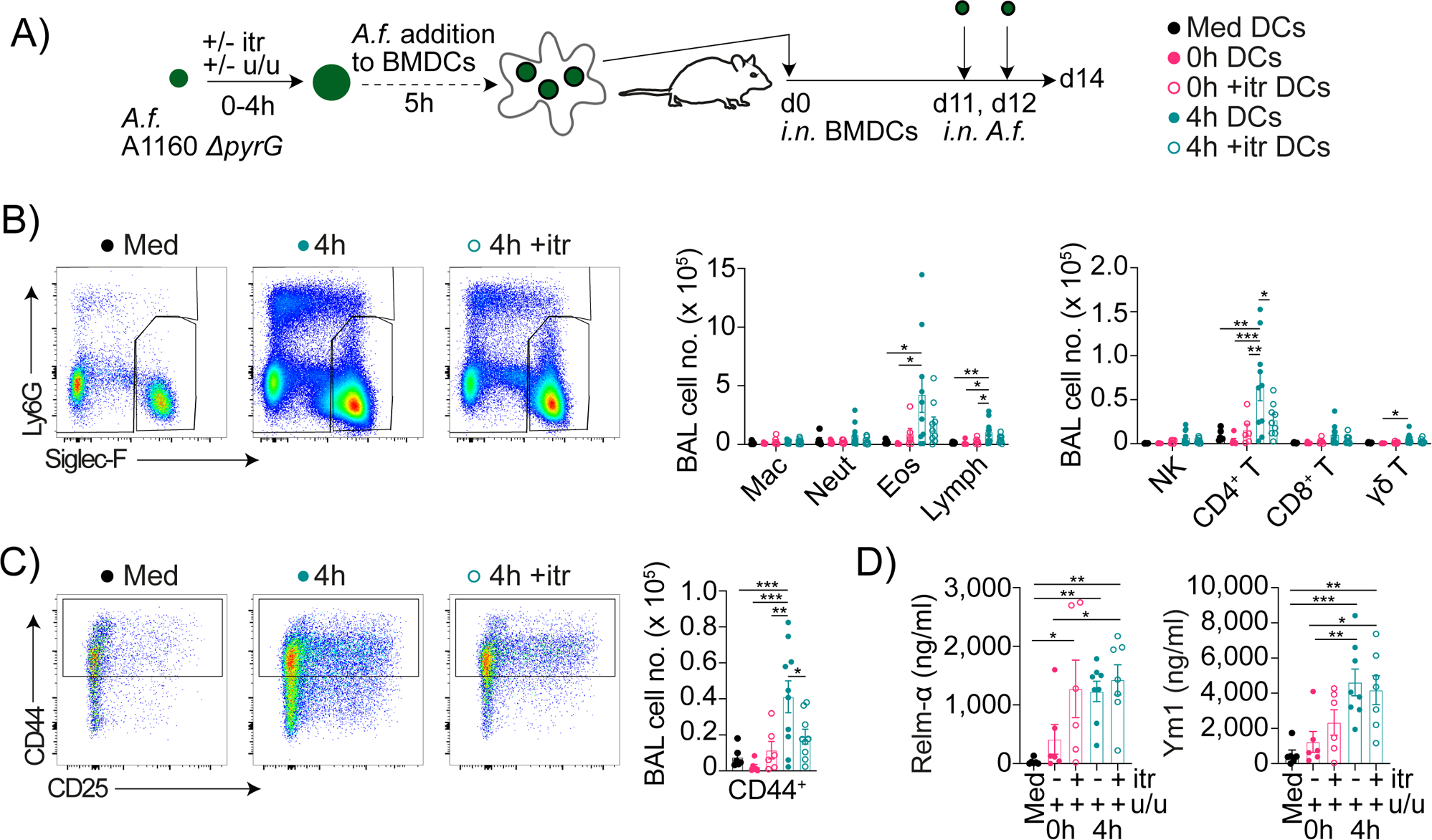
Spores treated with itraconazole reduces their capacity to elicit DC mediated fungal allergic airway inflammation. **A**) *ΔpyrG* spores were incubated with uridine and uracil (u/u) containing media in the presence or absence of itraconazole (itr) then growth was arrested by the removal of u/u media at 0 or 4 h. These were then incubated with DCs for 5h, prior to *i.n.* transfer of 50,000 DCs into C57BL/6 mice. Mice were challenged *i.n.* at d11 and 12 with 0.4 × 10 live CEA10 *A.f.* spores. **B**) Immune cell populations in BAL fluid were assessed by flow cytometry. The number of different cell populations isolated from BAL fluid of DC sensitised mice were assessed by flow cytometry. **C**) Representative flow cytometry plots of CD44^+^CD4^+^ T cells in BAL fluid. **D**) Secretory factors in BAL fluid were quantified by ELISA. Data are combined from two independent experiments (n=6-10 per group). Each data point is an individual mouse. Linear mixed effect modelling applied, with experimental repeat as a random effect variable, to compare multiple groups a post-hoc Tukeys HSD test was used. *=P <0.05,**=P <0.01, ***=P <0.001, ****=P<0.0001.

## Discussion

Using novel approaches to track and arrest differing stages of *A*.*f*. germination we have identified that isotropic spore growth is essential for DCs to elicit allergic inflammation after *A.f*. exposure. We have further identified that itraconazole, a crucial antifungal drug, can limit this process thereby reducing the ability of *A.f.* to elicit DC induction of allergic airway inflammation.

Sensitisation to *A.f.* triggers and exacerbates allergic airway inflammation that underpins asthma (17,29,42). Yet few studies have dissected the relative importance of early spore growth in triggering innate immune cells to elicit allergic inflammation. Previous studies have proposed that germination, with the breakdown of the hydrophobic rodlet/melanin layer that masks *A.f.* cell wall PAMPs e.g. β-glucan, is crucial to trigger innate cell activation and allergic inflammation (26,29). These findings were largely achieved using spore killing approaches (e.g. PFA) and we confirmed similar results, showing PFA-killed *A*.*f*. are less able to elicit allergic airway inflammation (Fig. 1) whilst germinated PFA-killed spores were more able to activate DCs (Supplementary Fig. 5). However, PFA-killing alters surface epitopes, disrupts spore metabolism and significantly changes dispersal patterns in the airways all of which may have caused the observed defect independent of disruption of spore germination (43,48). Our study overcomes these limitations, by utilising a mutant strain (*ΔpyrG*)(35,36) that that requires exogenous u/u to grow, we demonstrated a requirement for spore development to promote allergic airway inflammation, particularly in the BAL where dramatically reduced eosinophilia was observed after *i.n.* dosing with arrested (0h *ΔpyrG*) spores (Fig. 2). Surprisingly, *ΔpyrG* spores did induce minor lung inflammation, greater than the response elicited by PFA-killed spores, albeit to a much lower extent than responses caused with WT spores. Previous reports have shown that viable ungerminated spores are partially active, with low-level transcriptional activity (49). The mechanism(s) by which viable ungerminated spores can trigger low levels of inflammation is an important question for future work. In summary, these results highlight that breaking of dormancy and isotropic growth are crucial for the promotion of allergic airway inflammation.

We have recently identified a crucial role for DCs in mediating allergic inflammation in response to A.f. spores (17). Therefore, having established a requirement for spore swelling, we wanted to pinpoint if there was a precise *A.f.* morphotype which triggers DCs to induce allergic inflammation. Via novel use of the *ΔpyrG A.f.* strain with timed removal of u/u (Fig. 3), we discovered that the 3h viable early swollen *A.f.* spore stage, is the earliest stage able to induce DC activation and allergic priming (Fig. 3 & 4). To our knowledge, we are the first to demonstrate that an early-stage spore morphotype is crucial to generate allergic inflammation, earlier than previously thought as it is prior to the development of germlings and hyphal extension. Indeed, prior research on fungal allergy have focused predominantly on *A.f.* extracts and/or components revealed/secreted at the germling or hyphal stages that are not exposed at the spore stage. For example, research into β-glucan, a key mediator of antifungal responses and allergic inflammation, has focused on its exposure at the germling stage ∼ 7h, but recent work has suggested potential earlier expression (29,50). Furthermore, many of the previously characterised *A.f.* allergens are expressed/secreted predominantly at the hyphal stage, such as Asp f 5 and Asp f 13 (32,33,51,52), suggesting these may not be involved in initial spore mediated sensitisation. Therefore, this has caused us to be unaware of crucial mediators expressed by spores which trigger allergic disease. Recent evidence has emerged that the spore wall is more dynamic than previously thought with both resting and swollen spores (at 5h) expressing over 140 surface proteins, including low levels of β-glucan, and that earlier (2h) swollen *A.f.* spores able to secrete immunomodulatory proteases (34,50,53). Our novel approach of timed removal u/u to generate arrest *ΔpyrG* spore growth, could be crucial to elucidate immunogenic factors expressed at early stages of spore development, which have the potential to interact with the host immune cells.

To ascertain the potential of *A*.*f*. spores to elicit allergy we primarily focused on their interactions with DCs, due to their well-established role bridging innate and adaptive immunity to initiate allergic airway responses (12,14). Determining the ability of DCs to mediate type 2 inflammation *in vitro* is challenging as the mechanism(s) they utilise to mediate these responses are unclear(54). However, 3h spores increased DC expression of co-stimulatory molecules (CD80 and CD86) and pro-inflammatory cytokine prevalent during allergic inflammation (55–58). To confirm these changes resulted in allergic inflammation we utilised a DC transfer approach, which we have used previously with other stimuli (39,59). This showed 3h spore pulsed DCs induced a type 2 inflammatory response (including influxes of eosinophils and type 2 cytokine expressing CD4^+^ T cells) (Fig. 4) similar to responses observed following repeat *i.n. A.f.* exposure and demonstrates the importance of *in vivo* testing when confirming capability of DCs to induce type 2 polarisation. The precise process that DCs utilise to recognise spores and trigger downstream inflammation is unclear. However, *A.f.* and allergens such as HDM has been shown to signal via both Toll-Like receptor and C type lectin receptors on innate immune cells (60–62). Defining the precise spore morphotype that elicits DCs to mediate allergic inflammation provides an important platform to address this crucial question.

Despite evidence that antifungal drugs can be utilised as effective treatment strategies for SAFS and ABPA patients, the fungal morphotypes targeted in this intervention are unclear (45–47). Here, we identify a possible mechanism, in which antifungal treatment inhibits the early stages of spore swelling, which we have shown to be critical to activate DCs to prime fungal airway allergy (Fig. 6). Notably, these results could also explain the efficacy observed in clinical trials of itraconazole in severe asthma independent of fungal colonisation status, as our data indicate that spores would only need to remain in the airways for 3h to induce allergic inflammation (63). Furthermore, our data highlight that itraconazole treatment alone is not sufficient to completely arrest spore swelling and inhibit their ability to induce DC mediated allergic airway inflammation. Recent work has identified that combination therapy of antifungal agents can be more effective in treatment of disease and reduce the threat of drug resistance emerging (64). Therefore, with more therapeutics needed, high-throughput screening of compound libraries to identify candidates able to inhibit early spore swelling may be an effective strategy for discovery of novel drugs to treat fungal allergy (65).

Taken together, our work has highlighted the critical importance of the early stages of spore swelling in promoting allergic airway inflammation, with understanding the allergenic factors in 3h spores a key angle for future research. Additionally, our work extends understanding of the potential mechanism of action of a currently used drug (itraconazole) in fungal asthma, whilst also recommending that future therapeutics should be assessed for their ability to inhibit isotropic growth. Although our focus has been on *A.f.*, our findings are relevant for other allergenic filamentous fungi, including other *Aspergillus* species such as *A. flavus* and *A. niger*, with aspergillus sensitisation associated with increased asthma severity (66–69).

## Acknowledgements

We would like to thank current and previous members of the Manchester Fungal Infection Group, Lydia Becker Institute and MacDonald laboratory (University of Manchester) and MRC Centre for Medical Mycology (University of Exeter) for scientific discussions and some experimental assistance, and the University of Manchester and Exeter flow cytometry facilities. ELH was supported by a BBSRC CASE studentship (with GSK) (BB/P504543/1) and ASM by Lydia Becker core funding and the MRC (MR/W018748/1). PCC was supported with a University of Manchester Dean’s Prize Early Career Research Fellowship, Springboard Award (Academy of Medical Sciences, grant no. SBF002/1076). GV, DC and PCC were supported by Royal Society and Wellcome Trust Sir Henry Dale Fellowship (218550/Z/19/Z, awarded to PCC). GV, DC, GDB and PCC acknowledge funding from the MRC Centre for Medical Mycology at the University of Exeter (MR/N006364/2 and MR/V033417/1), the NIHR Exeter Biomedical Research Centre and the MRC Doctoral Training Grant MR/W502649/1. GDB acknowledges funding from the Wellcome Trust (217163). Additional work may have been undertaken by the University of Exeter Biological Services Unit. The views expressed are those of the author(s) and not necessarily those of the NIHR or the Department of Health and Social Care. SG was co-funded by the National Institute for Health Research (NIHR) Manchester Biomedical Research Centre (https://www.manchesterbrc.nihr.ac.uk/) and a National Centre for the Replacement, Refinement & Reduction of Animals in research (NC3Rs, https://www.nc3rs.org.uk/) Training Fellowship (NC/P002390/1), the Fungal Infection Trust and the Dowager Countess Eleanor Peel Trust.

## Supplementary Figure Legends

**Supplementary Figure 1. Flow cytometry gating strategy for the identification of immune cells in BAL** Representative flow cytometry plots showing gating for cells isolated from *A.f.* exposed BAL to identify macrophages (CD64^+^ MerTK^+^), eosinophils (MerTK^−^ SiglecF^+^), Neutrophils (Ly6G^+^ CD11b^+^), B cells (CD19^+^ MHCII^+^), γδ T cells (TCRγδ^+^), CD4^+^ and CD4^−^ T cells.

**Supplementary Figure 2. Flow cytometry gating strategy for the identification of cytokine expressing T cells from lung tissue.** Representative flow cytometry plots showing gating for cells isolated from naive lung tissue to identify CD4^+^ Foxp3^−^ T cells expressing various cytokines (IL-13, IL-17, IFNγ, IL-4, IL-5 and IL-10) after *ex vivo* stimulation.

**Supplementary Figure 3. Flow cytometry gating strategy for BMDC activation assessment.** Representative flow cytometry plots showing gating for BMDCs post 5h stimulation with spores, media or LPS. Costimulatory molecules and MHCII are used as activation markers - CD40, CD80, CD86.

**Supplementary Figure 4. Microscopic determination of spore size during germination.**A) Representative images of WT or *ΔpyrG* spores that were incubated in the presence or absence of uridine and uracil (u/u) with growth arrested by the removal of u/u media at the indicated timings (between 0 - 6h). Area of individual spores (100-1000 per group) was calculated using FIJI (70) using an in-house macro (Supplementary File 1) making use of the MorphoLibJ library (71). Briefly, images of spores from every condition and timepoint were segmented and measured for area per cell, per timepoint. To verify performance of the macro manual measurement of 30 0h and 30 6h spores was performed and compared to the macro, with over 95% agreement. B) *ΔpyrG A.f.* spores (A1160) were incubated at 37°C for 6h in the presence of uridine and uracil (u/u), u/u media was removed, and spores were left at 4°C for 14 days, on day 14 media with u/u was added to the spores and they were left overnight at 37°C. The next day, hyphal structures were imaged using a light microscopy with a 20x phase contrast objective lens.

**Supplementary Figure 5. Spore swelling, but not viability, is required for DC activation.** DCs were incubated for 5h with either live *A.f.* spores (A1160) (MOI 1:1, 1:5 or 1:10) or PFA-killed spores (MOI 1:1 or 1:5). PFA killed spores were either generated from uncultured spores or spores that had been incubated at 37°c for 4 hours. **A**) DC expression of co-stimulatory molecules and MHCII was analysed via flow cytometry. **B**) DC cytokine secretion was measured via ELISA. Each data point is an independent experiment (n = 5). Mixed effect modelling applied, with post-hoc multiple comparisons between groups. Statistical comparisons to LPS positive control are not shown. *=P <0.05, **=P <0.01, ***=P <0.001.

