## Supplementary figures and images for "*Aspergillus*-mediated allergic airway inflammation is triggered by dendritic cell recognition of a defined spore morphotype, a process that can be targeted via antifungal therapeutics"

### Supplementary Figure 1

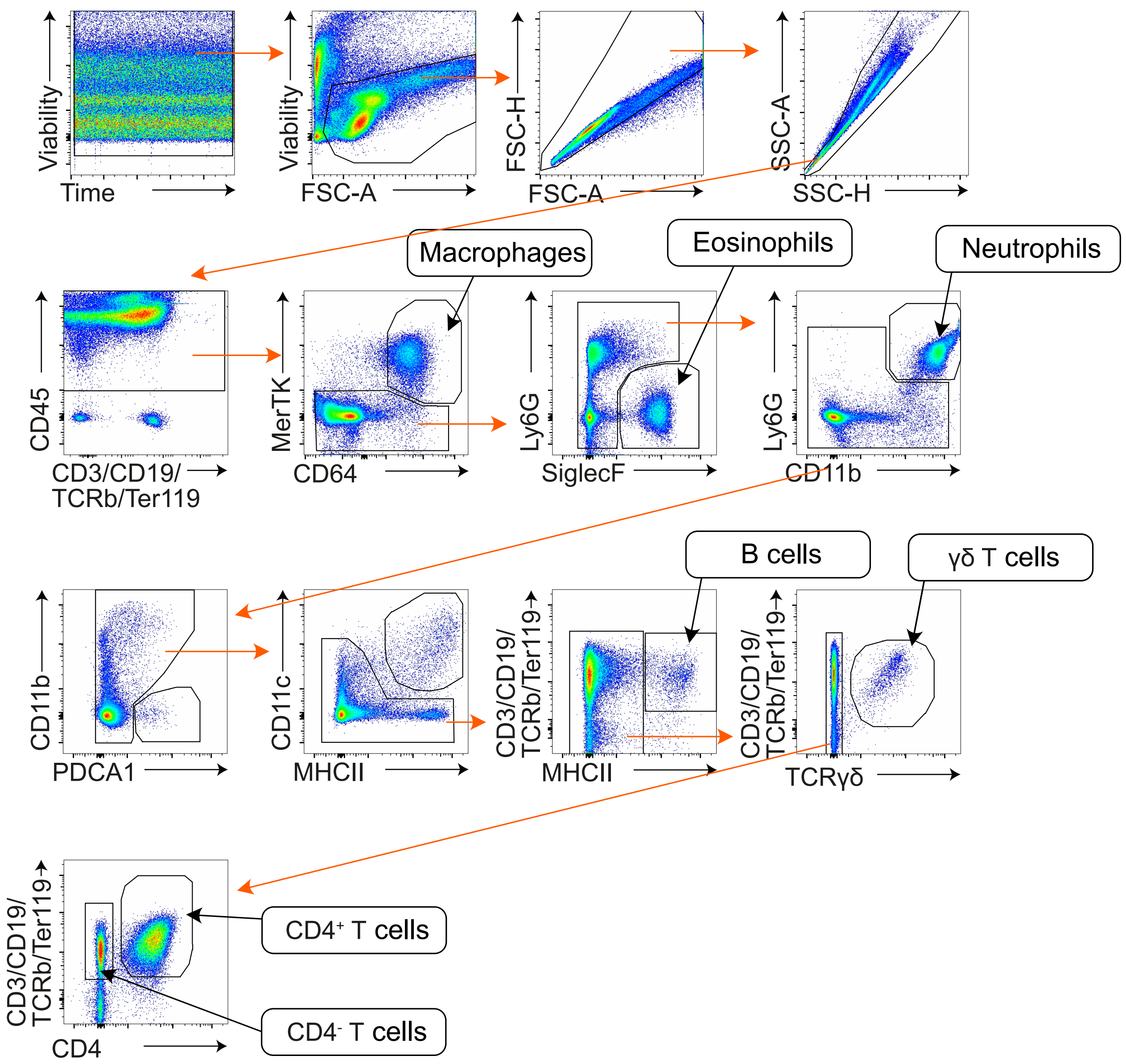


**Supplementary Figure 1**

### Supplementary Figure 2

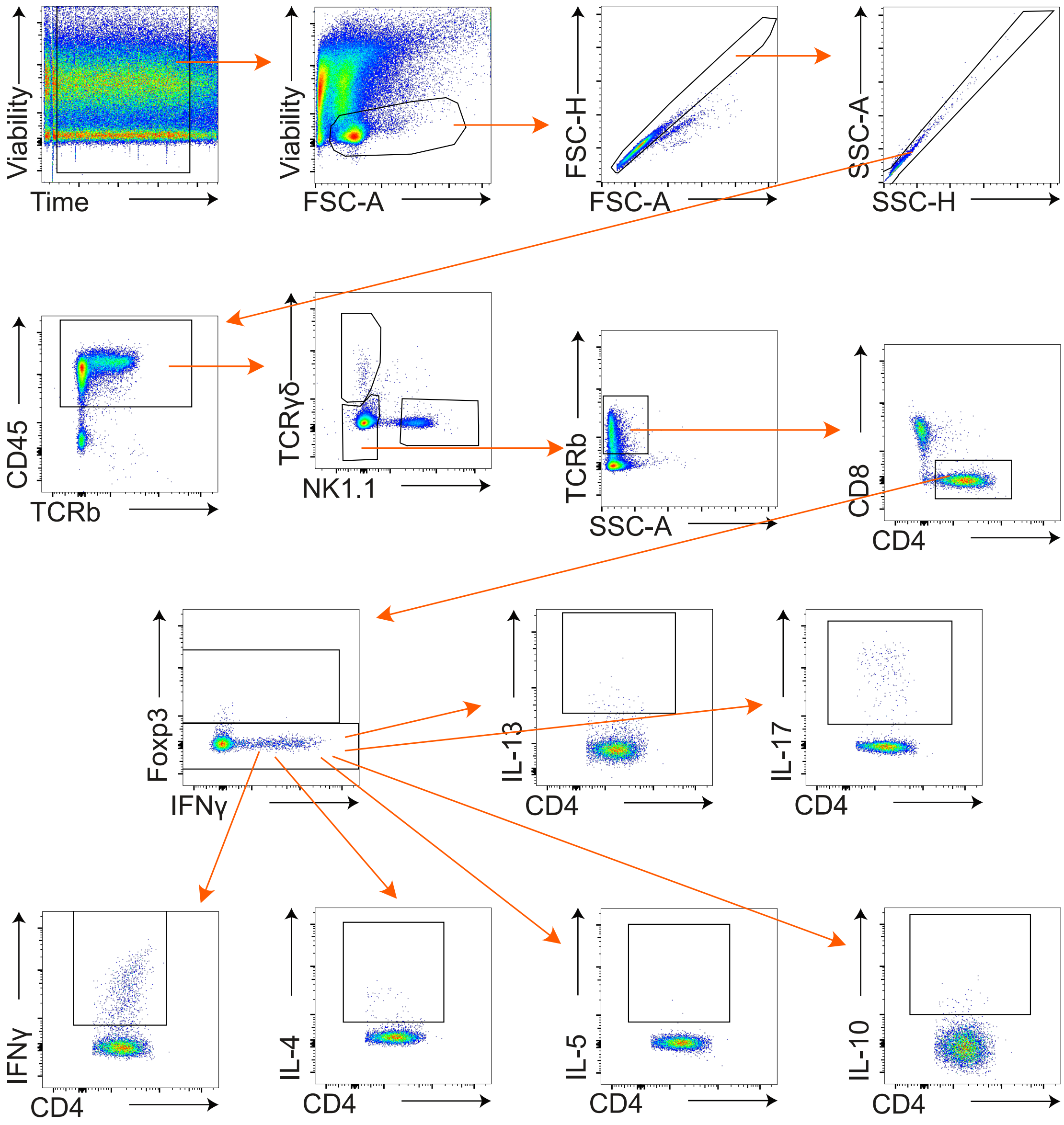


**Supplementary Figure 2**

### Supplementary Figure 3

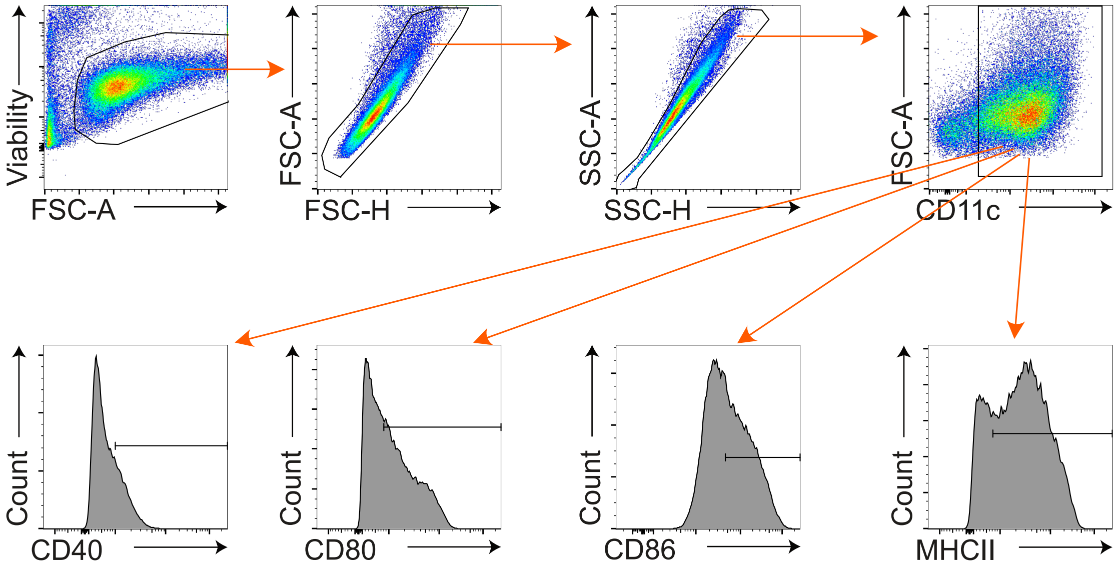


**Supplementary Figure 3**

### Supplementary Figure 4

A)

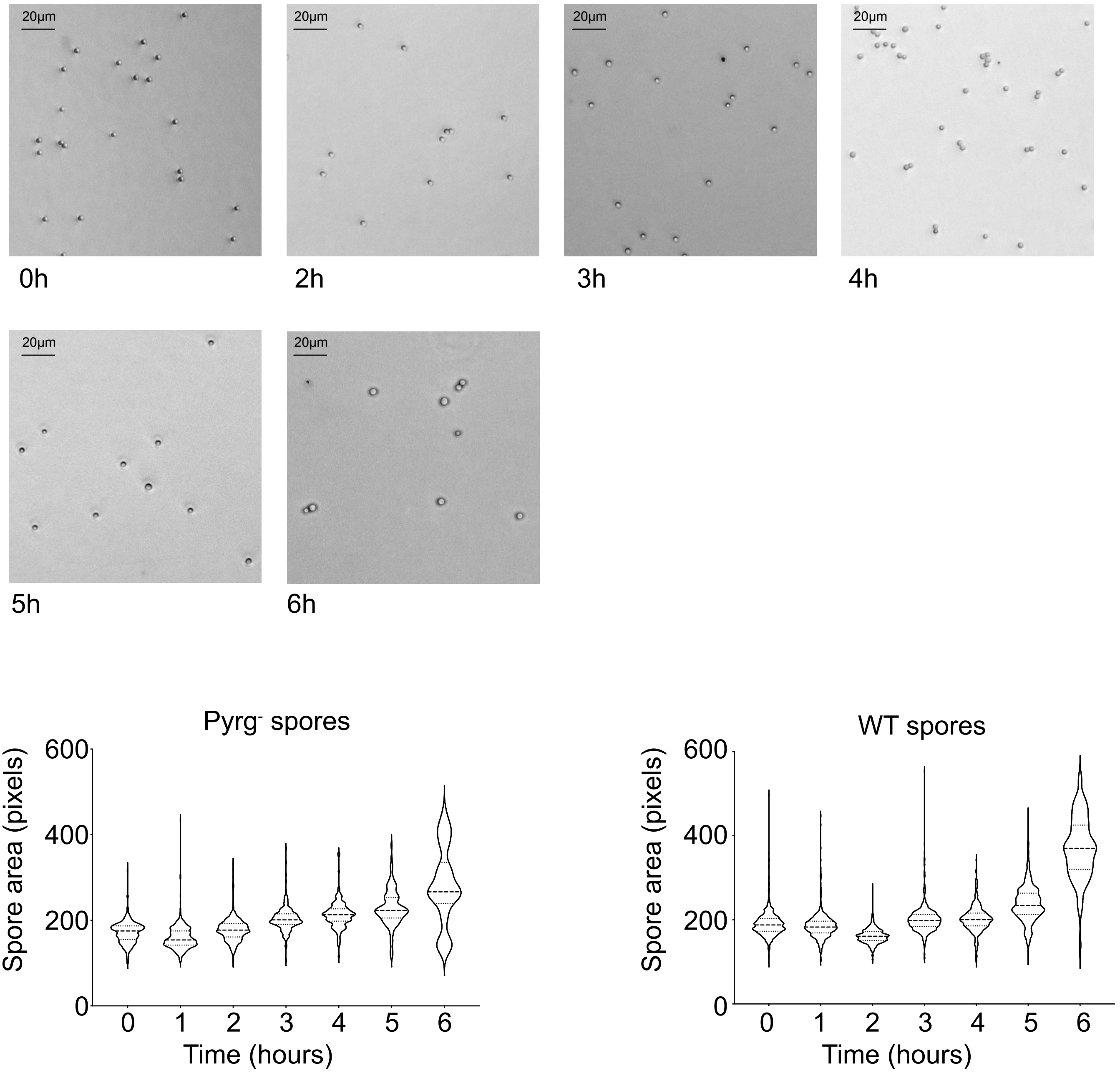


B)

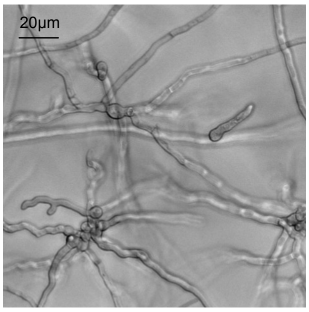


Pyrg$^-$ spores
Post 24hrs in U/U media

**Supplementary Figure 4**

### Supplementary Figure 5

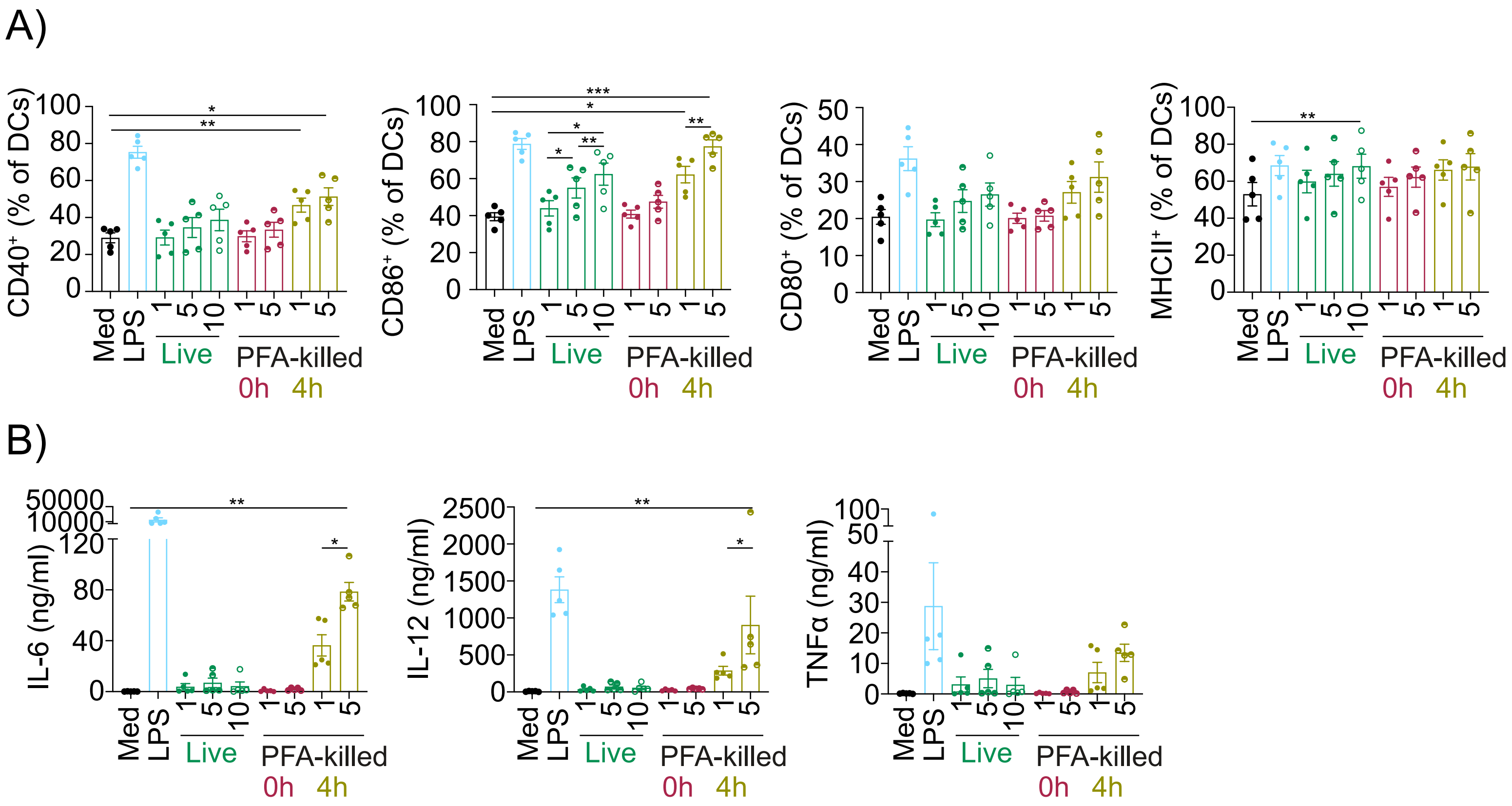


**Supplementary Figure 5**
